# Single-nuclei Transcriptome of Human AT Reveals Metabolically Distinct Depot-Specific Adipose Progenitor Subpopulations

**DOI:** 10.1101/2022.06.29.496888

**Authors:** Clarissa Strieder-Barboza, Carmen G. Flesher, Lynn M. Geletka, Jennifer B. Delproposto, Tad Eichler, Olukemi Akinleye, Alexander Ky, Anne P. Ehlers, Robert W. O’Rourke, Carey N. Lumeng

## Abstract

Single-cell and single-nuclei RNA sequencing data (scRNAseq and snRNAseq, respectively) have revealed substantial heterogeneity in the AT (AT) cellular landscape in rodents and humans depending on depot and disease status. We used snRNAseq to characterize the cellular landscape of human visceral (VAT) and subcutaneous AT (SAT) samples from lean subjects and subjects with obesity. We identified multiple cell types in the AT cellular repertoire, including three major AT stromal cell (ASC) subpopulations, multiple types of adipocyte (ADIPO) populations that retain properties similar to ASC, endothelial cell (EC), T-cell, and macrophage (MAC) populations that are in concordance and expand upon other published datasets. ADIP and EC are more prominent in SAT compared to VAT which has a higher proportion of ASC. Of two dominant ASC subpopulations, one (inflammatory-mesothelial, IM- ASC) is present in VAT, but absent in SAT, while the other (fibroadipogenic, FA-ASC) is present in both VAT and SAT. Both populations retain their properties in *in vitro* culture and have adipogenic capacity with different metabolic features. Informatic and *in situ* studies support ADIP derived from IM- and FA- ASC are found in human VAT. We also identified a wide range of EC subtypes in human AT with features of lymphatic, venous, and arterial EC, with identification of a *PRDM16* expression EC population with features of an EC progenitor. Immune cell populations match recent experimental validation of lipid activated macrophage (LAM) phenotypes, TIM4 macrophages, and a prominent population of MRC1/CD206^+^ resident AT macrophages with gene expression signatures related to glucocorticoid activation. Overall, our study demonstrates depot-specific cellular diversity in human VAT and SAT in which distinct ASC subpopulations may differently contribute to AT dysfunction in obesity. Also, our results highlight an unprecedented EC heterogeneity suggesting AT EC as highly specialized cells and potentially important regulators of depot-specific functions.

**METHODS STATEMENT:** Human subjects provided informed consent and were enrolled with approval from Institutional Review Boards at the University of Michigan and Veterans Affairs Ann Arbor Healthcare System. Enrollment, consent, and all aspects of human subject research were carried out in accordance with the Belmont Report from the National Research Act of 1974, and the Declaration of Helsinki set forth by the World Medical Association. This manuscript contains no human participants’ names or other HIPAA identifiers.

## Introduction

Single-cell RNA sequencing (scRNAseq) data in mice and humans confirm significant heterogeneity of the AT cellular landscape and identify multiple functionally distinct populations of adipocyte progenitor cells (preadipocytes), immune cells, and other cell types. ^1–6^ This cellular diversity has important implications for an understanding of obesity and metabolic disease pathophysiology and development of translational therapy. scRNAseq methodology has numerous limitations for analysis of AT, including the requirement for collagenase digestion of tissue, which may alter the transcriptome and eliminate cell subpopulations, the inability to capture large, fragile mature adipocytes which are disrupted by microfluidic sorting, and the inability to study banked, frozen samples. To address these limitations, several research groups have performed single-nuclei RNA sequencing (snRNAseq) in AT, permitting isolation of nuclei from all cells within AT, including adipocytes, from fresh and frozen untreated tissue samples.^7–11^

Depot-specific differences in AT stromal cell (ASC) and mature adipocyte metabolic phenotypes are well-established in human and mice models. Omental VAT adipocytes, relative to abdominal SAT adipocytes, are smaller in size and exhibit decreased insulin sensitivity and adipogenic capacity, and increased lipolytic response, features that correlate with systemic hyperglycemia and hyperlipidemia.^12–,15^ The correlation between ASC/adipocyte dysfunction and systemic metabolic disease in humans supports investigation of ASC and adipocytes subtypes as high yield targets in deciphering AT depot-specific function. The majority of prior human research in this area evaluates adipocytes differentiated from stromal vascular fraction (SVF) isolated from AT. We now know that this approach lacks rigor, as scRNAseq data from mice and humans demonstrate substantial heterogeneity in ASC and adipocytes.^5, 6, 16, 17^

Our goal was to apply snRNAseq analysis to banked frozen AT and describe the complete transcriptome of human VAT and SAT. We studied VAT and SAT samples from a cohort of lean human subjects and subjects with obesity. We identified an snRNAseq landscape that includes all expected subpopulations of cell types, and multiple subpopulations of ASC and mature adipocytes (ADIP) that correspond to those described by others.^2, 7, 18^ We also identified transcriptionally and metabolically distinct populations of ASC with depot-specific differences in frequency. These findings clarify the evolving understanding of human AT cellular composition and suggest possible mechanisms underlying depot-specific differences in human AT biology.

## Results

### Single nuclear resolution transcriptomics of human AT expands cell type diversity

We adapted published methods to perform snRNAseq in eleven AT samples derived from eight individuals (Table 1)^19^. After filtering, the dataset included 36,588 nuclei isolated from human VAT and SAT of lean human subjects and subjects with obesity (Figure 1A). Manual annotation of the initial cluster analysis identified nuclei expressing signature genes consistent with adipocytes (ADIPO; *ADIPOQ, LEP*), macrophages (MAC; *MERTK*), T-cells (TCELL; *CD69*), endothelial cells (EC; *VWF*), pericytes (PC; *NOTCH3*), and lymphatic endothelial cells (LEC; *MMRN1*). (Figure 1B-D, Table S1). In addition, an EC population with progenitor markers was also identified with high expression of *PRDM16* (EC-P), a marker of adipogenic cells^20^. Three discrete subpopulations of ASC were identified based on expression of *PDGFRA* and *PDGFRB*. To complement our manual annotation, we calculated expression scores for the top 100 genes from a human GTEX snRNAseq dataset from breast and found strong concordance with their cellular computational annotations and our major cell populations.^8^ (Figure 1C Vijay *et al.* has previously published a scRNAseq data set from human VAT and SAT^1^.

**Figure 1.**
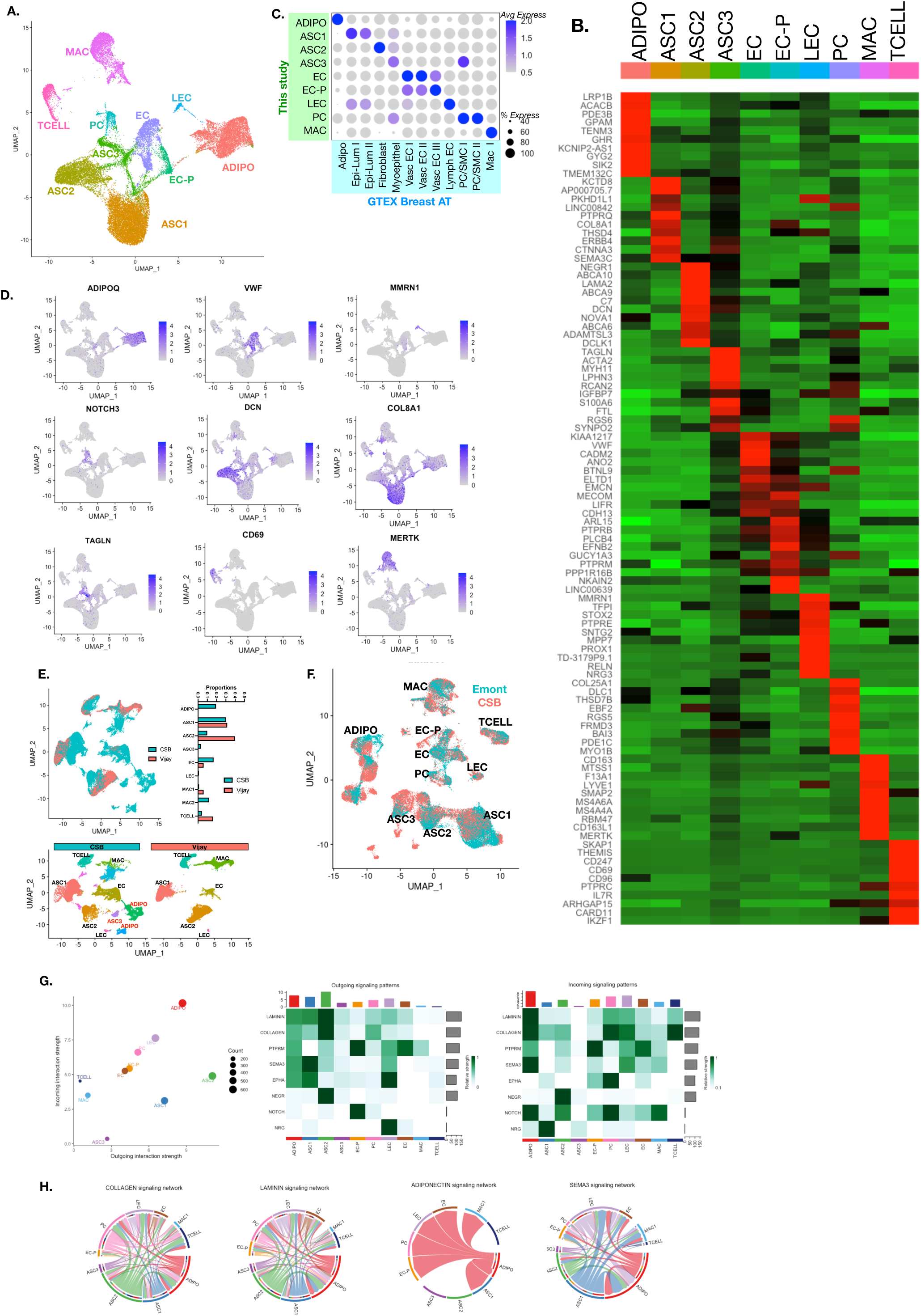
snRNAseq identifies distinct cell subpopulations in human AT. (A) UMAP plot of nuclei sub-populations in combined omental visceral (VAT) and abdominal subcutaneous (SAT) AT samples from human subjects. Cell populations were classified as adipose stromal cells (ASC), mature adipocytes (ADIPO), endothelial cells (EC), lymphatic endothelial cells (LEC), endothelial cell progenitors (EC-P), pericytes (PC), macrophages (MAC), and T-cells (TCELL). (B) Heatmap of top DE genes in cell subtypes identified. (C) Module scores computed for cells in our study based on DEG identified in the GTEX Breast AT cell types. (D) UMAP plots of signature genes for ADIPO (*ADIPOQ*), EC (*VWF*), LEC (*MNRN1*), PC (*NOTCH3*), ASC (*DCN, COL8A1, TAGLN*), TCELL (*CD69*) and MAC (*MERTK*). (E) Integrated analysis of nuclear cell types in our dataset compared with Vijay et al scRNAseq data. Cell type proportions in Vijay et al (red) and this dataset (blue). (F) Integrated analysis of nuclear cell types in this dataset and Emont et al. (G) Cellchat analysis of major populations of snRNAseq dataset. Plot of outgoing and incoming interactions signal strengths for cell types and heat map of prominent signaling pathways. (H) Circle plots of most prominent cell-cell communication pathways identified in analysis of interactions in the full dataset.

**Table 1.** Patient Demographics for samples.

| Patient | Tissue studied | Sex | Age | BMI | DM status | HbA1c | Ethnicity | HTN | OSA | HL | Insulin | Metformin | Statin |
| --- | --- | --- | --- | --- | --- | --- | --- | --- | --- | --- | --- | --- | --- |
| 1 | VAT, SAT, | F | 52 | 43 | DM | 7.9 | Caucasian | Y | N | Y | Y | Y | N |
| 2 | VAT, SAT, | M | 46 | 28 | NDM | NA | Caucasian | Y | N | N | N | N | N |
| 3 | VAT, SAT, | F | 34 | 63 | NDM | 5.5 | Caucasian | N | N | N | N | N | N |
| 4 | VAT, SAT, | M | 64 | 26 | NDM | NA | Caucasian | Y | N | N | N | N | N |
| 5 | VAT | F | 47 | 24 | NDM | 4.8 | Caucasian | N | N | N | N | N | N |
| 6 | VAT, SAT, V-ASC | F | 36 | 43 | NDM | 5.7 | Caucasian | N | Y | N | Y | N | N |
| 7 | V-ASC | F | 32 | 45 | PRE-DM | 5.9 | Black | N | N | N | N | N | Y |
| 8 | V-ASC | F | 39 | 46 | DM | 5.6 | Caucasian | N | N | N | N | N | Y |
| 9 | V-ASC | F | 30 | 43 | DM | 5.8 | N/A | N | Y | N | N | N | Y |

Tissue studied: VAT: visceral AT studied with snRNASeq; SAT: subcutaneous AT studied with snRNASeq; V-ASC visceral AT -derived ASC studied with scRNASeq
DM, PRE-DM, NDM defined by ADA HbA1c criteria off hypoglycemic medications and by medication use
HTN: hypertension; OSA: obstructive sleep apnea; HL: hyperlipidemia
HbA1c within 6 months of surgery/tissue collection, while treated with DM medication

Integration of our dataset (“CSB”) with Vijay *et al.’s* showed overlapping cell types for ASC1, ASC2, EC, LEC, MAC, and TCELL (Figure 1E). Our snRNAseq dataset captured numerous cell types that were not readily identified in the Vijay dataset. As expected, ADIPO were not represented in Vijay *et al.’s* study, as they performed scRNAseq of SVF. In addition, ASC3 identified in our dataset was not seen in Vijay’s study and the proportion of EC were significantly underrepresented in their dataset. These observations are consistent with concerns of inefficient cell capture of all AT cell types with traditional collagenase-based cell dissociation methods and the loss of lipid laden cell types that may be closely associated with buoyant adipocytes. ^21^ Recently, Emont *et al* published a large snRNAseq dataset from human and mouse AT. Integration of our dataset with Emont’s demonstrated close alignment of our cell types and those identified by their work (Figure 1F).

To understand potential cell-cell paracrine interactions between all cell types in our snRNAseq dataset, we performed analysis with CellPhoneDB on all cell types as a whole^22^ (Figure 1G-H). Ranking cell types based on incoming and outgoing interaction strengths showed that ADIPO had the highest interactions with other cell types. ASC2 generated more outgoing signals than incoming signals. PC and EC cell types had similar proportions of incoming and outgoing interactions and were clustered below ADIPO. The most prominent outgoing signaling patterns related to laminin and collagen signaling with highest signals from ASC2. ADIPO were enriched for laminin, collagen, and neuronal signaling pathways (P*TPRM, SEMA3, EPHA*). Neuregulin and ephrin outgoing signals were enriched in LEC. *NOTCH* and *PTPRM* (Protein Tyrosine Phosphatase Receptor Type M) outgoing signals were enriched in EC-P cells. In terms of dominant incoming responses, ADIPO had the highest incoming signals from laminin, collagen, *NOTCH* and *SEMA3* signaling pathways. Evaluation of canonical adipocyte signaling pathways involving adiponectin identified multiple signals between adipocytes and multiple stromal cells including EC, LEC, PC, EC-P, MAC, and ASC1 (Figure 1H).

### Depot specific differences in AT cellular composition

We focused our initial analysis on differences in cellular composition and gene expression between human SAT and VAT (Figure 2A). All major cell types were represented in both SAT and VAT (Figure 2B). Comparing proportions of cell types present in each depot, there was a significant increase in ADIPO in SAT compared to VAT. Notably, our snRNAseq data demonstrates that SAT ADIPO nuclei compose ∼30% of nuclei in SAT but only ∼15% of nuclei in VAT. In addition, EC and EC-P populations were enriched in SAT compared to VAT, consistent with reports of increased angiogenic potential in SAT compared to VAT^23^. To experimentally validate these observations, flow cytometry was used to evaluate SVF cells in paired VAT and SAT samples from obese patients undergoing bariatric surgery (Figure 2C), which demonstrated an enrichment of EC (CD31^+^CD45^-^) in SAT and a decrease in ASC (CD34^+^CD45^-^) in SAT compared to VAT.

**Figure 2.**
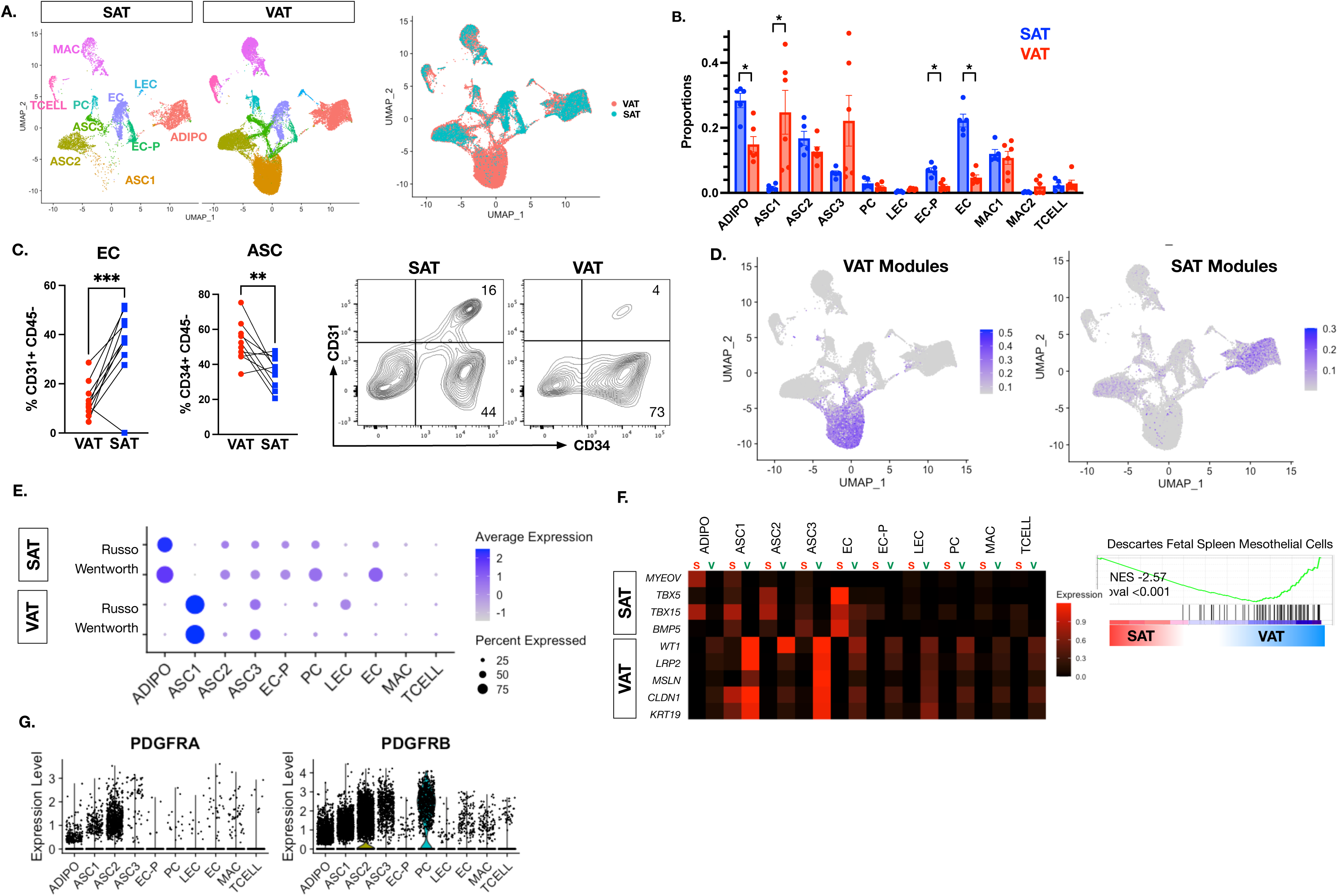
Depot specific differences in cell types. (A) UMAP plots of VAT and SAT specific cells. (B) Cell type proportions in SAT (blue) and VAT (red) samples (n=11). T-test was used to compare SAT vs. VAT cell frequencies and were considered significant when P<0.05. (C) Quantitation of EC (CD31^+^CD45^-^) and ASC (CD31^4^CD45^-^) in matched VAT and SAT from obese patients by flow cytometry (n=10). Representative flow cytometry plots shown. Paired t-test was used to compare SAT *vs.* VAT EC and ASC frequencies and were considered significant when P<0.05. (D) FeaturePlot of Expression scores for core VAT and SAT gene modules of the top 50 DE genes between depots based on Russo et al ^24^. (E) Dot plot of expression scores for gene modules for DE genes between SAT and VAT from Russo et al ^24^ and Wentworth et al. ^25^. (F) Heatmap of cell type specific expression patterns of the top DE genes in SAT (top) and VAT (bottom). Pathway analysis (GSEA) of mesothelial lineage genes. (G) Expression plots of *PDGFRA* and *PDGFRB* expression in our snRNAseq dataset.

The most prominent quantitative difference between depots was a specific enrichment of ASC1 in VAT, but not in SAT. Module scores were calculated for the top DE genes between VAT and SAT (Figure 2D). Scores for VAT prevalent genes were highest in the VAT-specific ASC1 population indicating that ASC1 genes dominate depot differences from bulk RNAseq analysis. SAT enriched gene modules were primarily concentrated in ADIPO and ASC2 populations. To assess if bulk RNAseq datasets comparing depots were attributable to specific cell types, we evaluated the top DE genes from our previously published bulk RNAseq analysis from obese human VAT and SAT (‘Russo’)^24^ and another comparison of VAT and SAT bulk RNAseq from Wentworth *et al* ^25^ (Figure 2E). Depot specific DE genes were used to calculate composite expression scores on a per cell basis. The SAT enriched genes from the bulk RNAseq data can be predominantly attributed to SAT adipocyte gene expression, potentially due to the ∼2-fold increase in adipocytes in SAT compared to VAT as a proportion of all cells. DE genes identified as VAT specific were derived primarily from ASC1 due to the absence of this cell type from SAT.

Evaluation of the top DE genes further supported cell type specific gene expression that drives difference in bulk RNAseq analysis between SAT and VAT (Figure 2F). For SAT specific genes, *MYEOV* is specifically enriched in SAT ADIPO cells compared to VAT ADIPO. *TBX15* is enriched in multiple SAT cell types including ADIPO, ASC1, ASC2, and EC. *TBX5* is prominent in ASC2 and EC from SAT with low expression in VAT. For the top DE genes in VAT, most were enriched in ASC1 and ASC3 cells. Many of these are markers of mesothelial-like cells such as *WT1*, *MSLN*, *CLDN1*, and *KRT19 (*Figure 2F), which is consistent with Pathway analysis showing enrichment for mesothelial cell pathways in VAT compared to SAT. We noted that the function of mesothelial-like cells in human AT remains unresolved and most prior single-cell studies from mice and humans do not differentiate mesothelial cells from other ASC types. Overlap between mesothelial cells and ASC is unresolved. Human mesothelial cells express *PDGF* and *PDGFRA* ^26, 27^, therefore, differentiation of mesothelial-like cells and other ASC types by *PDGFRA* expression may not be sufficient. Consistent with this, ASC1, ASC2, and ASC3 were all found to have prominent expression of both PDGRFA and PDGFRB (Figure 2G).

### Depot specific differences in adipose stromal cells (ASC)

Sub-analysis of ASC subsets resolved three major subpopulations (Figure 3A, Table S2). A minor population had high expression of smooth muscle genes that overlap with ICAM1 ASC (SM-ASC) that included *TAGLN* and *MYH11* ^2^ (Figure 3B). We observed a striking difference between VAT and SAT in ASC1-derived cells which we term “inflammatory mesothelial-like ASC” (IM-ASC) that are present in VAT, but absent in SAT. Signaling pathway enrichment analysis of IM-ASC specific genes with SPEED2 ^28^ identified enrichment in multiple inflammatory gene pathways including *IL-1, TLR, TNF,* and hypoxia signaling to support a pro-inflammatory signature (Figure 3C). Pathway analysis (GSEA) comparing DE genes enriched in IM-ASC compared to FA-ASC demonstrated increased inflammatory response pathways, TNFA signaling, and TGFβ signaling (Figure 3D, Table S3). IM-ASC specific inflammatory genes included *IL16* and *IL18* previously implicated in metabolic disease ^29–32^ suggesting that IM-ASC-derived *IL-18* and *IL-16* may contribute to VAT dysfunction.

**Figure 3.**
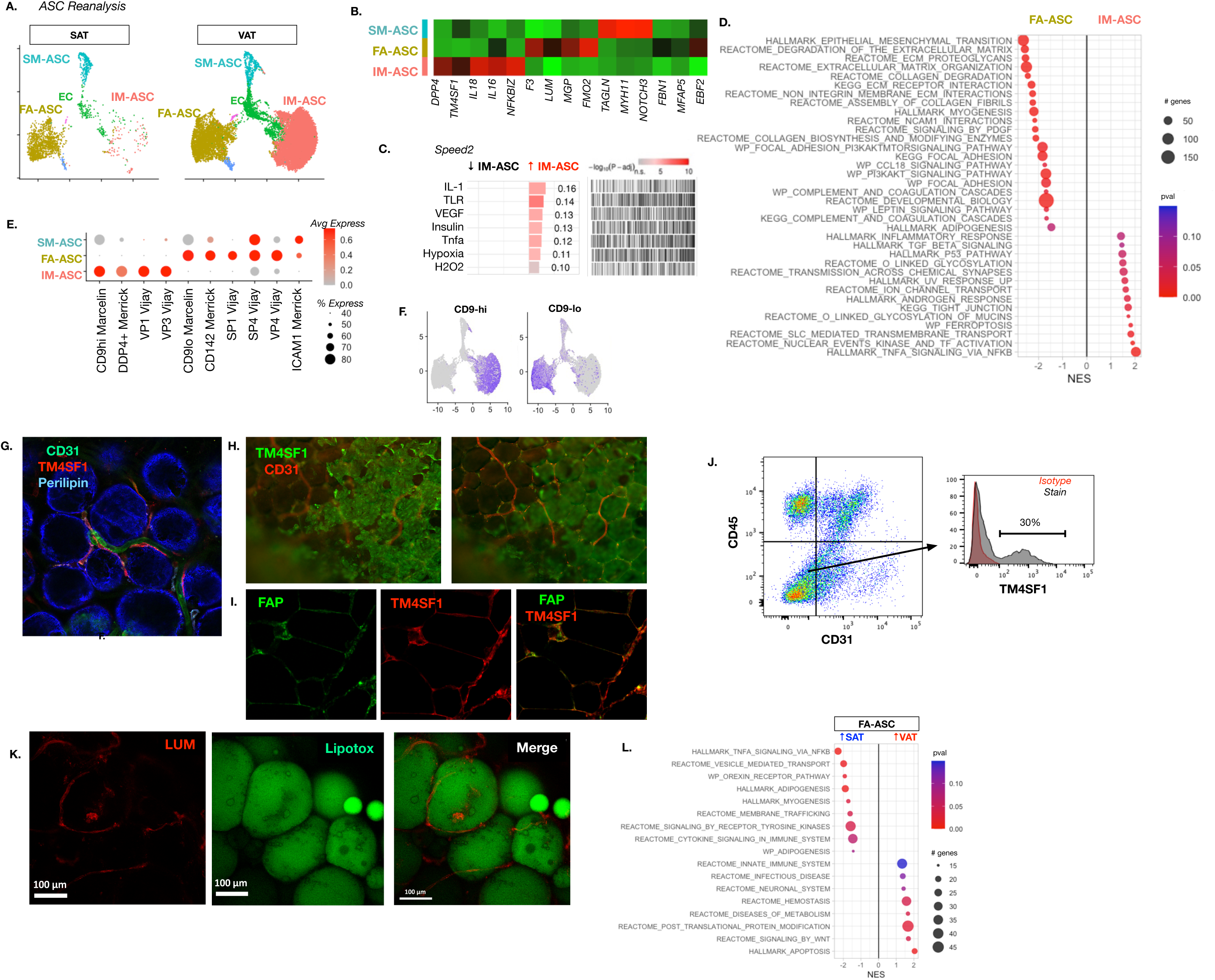
Depot specific differences in adipose stromal cells (ASC). (A) UMAP plots of subset analysis of ASC1-3 in dataset split by depot specific cells. (B) Heatmap of hallmark genes for primary ASC populations identified in subset analysis. (C) SPEED2 analysis of IM-ASC DE genes identify inflammatory signaling regulator pathways. (D) Pathway analysis (GSEA) of FA-ASC vs IM-ASC. (E) Dot plot of expression values of module scores for ASC identified in prior published studies from human AT. (F) Feature plot of gene module for CD9^hi^ and CD9^lo^ ASC identified in Marcelin et al ^18^. (G) Immunofluorescence staining of TM4SF1 (IM-ASC marker) in perivascular niche in human VAT (adipocyte marker perilipin in blue) in proximity to endothelial cells (CD31^+^). (H) Localization of TM4SF1^+^ cells on surface of whole mount VAT biopsy (left). Lower focal plane identifies sub-surface of TM4SF1^+^ cells in perivascular niche (right). (I) Immunofluorescence localization of TM4SF1^+^FAP^-^ cells on the surface of FPPE sections from human VAT distinct from TM4SF1^+^FAP^+^ cells below the cell surface using Opal. (J) Flow cytometry analysis and validation of TM4SF1^+^ (IM-ASC) and TM4SF1^-^ (FA-ASC) ASC in VAT SVF cells (CD31^-^CD45^-^). (K) Pathway analysis (GSEA) of DE genes between SAT and VAT in IM-ASC. (L) Immunofluorescence localization of LUM^+^ (FA-ASC) in VAT human AT biopsies by confocal microscopy.

To compare ASC identified here with other studies, we used the hallmark gene signatures of published human ASC types to compute expression scores for each ASC type (Figure 3E). IM-ASC have a gene expression signature that is enriched for genes identified in DPP4^+^ ASC described by *Merrick et al.* ^2^, mesothelial-like cells (MLC) by *Hepler et al*. ^5^, MSLN^+^ VP1/VP3 of *Vijay et al.* ^1^, and CD9^hi^ ASC described by *Marcelin et al.* ^18^ (Figure 3E-F). Our observations of VAT-specificity for IM- ASC agree with *Vijay et al.* for VP1/VP3 shown to have high mitochondrial activity. CD9^hi^ ASC described by *Marcelin et al.* were associated with VAT fibrosis and insulin resistance and overlap with our IM-ASC subpopulation. In mice, DPP4^+^ interstitial progenitor cells can form adipocytes in all depots but are induced to make adipocytes only in gonadal AT with obesity.^17^

We used immunofluorescence (IF) microscopy with the IM-ASC specific marker TM4SF1 (transmembrane 4 L six family member 1), a surface tetraspanin to identify the location of IM-ASC. TM4SF1^+^CD31^-^ IM-ASC were identified in the perivascular niche, distinct from TM4SF1^+^CD31^+^ EC (Figure 3G). IF of whole mount AT also identified TM4SF1^+^CD31^-^ mesothelial-like cells on the surface of human VAT. (Figure 3H) Importantly, we also found TM4SF1^+^CD31^-^ cells deeper in the strom. IF of AT sections co-stained with the ASC specific marker FAP (fibroblast activation protein) further supported TM4SF1^+^FAP^+^ IM-ASC found deeper in the stroma and TM4SF1^+^FAP^-^ cells on the surface lining of AT consistent with mesothelial-like cells (Figure 3I). Complementary flow cytometry analysis demonstrated CD31^-^CD45^-^TM4SF1^+^ in VAT (Figure 3J).

We term the prominent ASC subtype present in both depots (ASC2) “fibro-adipogenic ASC” (FA- ASC) based on enrichment in gene pathways related to AT extracellular matrix (ECM) remodeling, ECM organization, and collagen production. FA-ASC are enriched for genes identified in CD9^lo^ ASC described by *Marcelin et al.*, CD142^+^ ASC described by *Merrick et al.*, and CFD^+^ SP1/SP4/VP4 populations of *Vijay et al.* that correlate with DM. IF identified cells positive for the FA-ASC specific protein lumican (LUM) in whole mount VAT (Figure 3K).

We compared VAT and SAT FA-ASC to identify depot-specific qualitative differences in gene expression within the FA-ASC type. Evaluation of DE genes between SAT and VAT FA-ASC with Pathway analysis (GSEA) showed that SAT FA-ASC are enriched for adipogenesis pathways (*CD36, PPARG, ENPP2*), while VAT FA-ASC were enriched Wnt signaling, metabolic disease, and ECM organization genes (Figure 3L). These data demonstrate that FA-ASC in VAT, relative to SAT, have an accentuated fibroblast phenotype that may contribute to depot differences in AT and systemic metabolism.

### Adipocytes derived from FA-ASC and IM-ASC manifest distinct metabolic phenotypes in vitro

IF of cultured SVF cells from human VAT identified distinct TM4SF1^+^ IM-ASC and LUM^+^ FA- ASC (Figure 4A). *In vitro* differentiated adipocytes from SAT were composed of largely LUM^+^ adipocytes, consistent with SAT specificity of FA-ASC (Figure 4B). In contrast*, in vitro* differentiated VAT preadipocytes generated LUM^+^ and LUM^-^ adipocytes, suggesting that both IM-ASC and FA-ASC can differentiate into adipocytes *in vitro*. To independently evaluate the adipogenic potential of TM4SF1^+^ IM-ASC and LUM^+^TM4SF1^-^ FA-ASC, FACS was used to isolate the two ASC subtypes from primary VAT from obese subjects. qRTPCR confirmed enrichment of *TM4SF1* expression in TM4SF1^+^ IM-ASC and *LUM* in TM4SF1^-^ FA-ASC (Figure 4C). VAT purified IM- and FA-ASC were expanded and subjected to adipogenic differentiation. Adipocytes derived from FA-ASC, compared to adipocytes derived from IM-ASC or bulk ASC, manifested increased expression of adipogenic genes (*PPARG, ADIPOQ, PLIN, ATGL*) (Figure 4D), and increased adipogenesis based on lipid accumulation (Lipidtox) – a phenotype that was observed even in minimal differentiation media (without TZD or INS) (Figure 4E). FA-ASC derived adipocytes also showed increased lipolytic responses compared to adipocytes derived from IM-ASC (Figure 4F). These data demonstrate that adipocytes derived from FA-ASC have enhanced adipogenic and lipolytic capacity compared to IM-ASC derived adipocytes.

**Figure 4.**
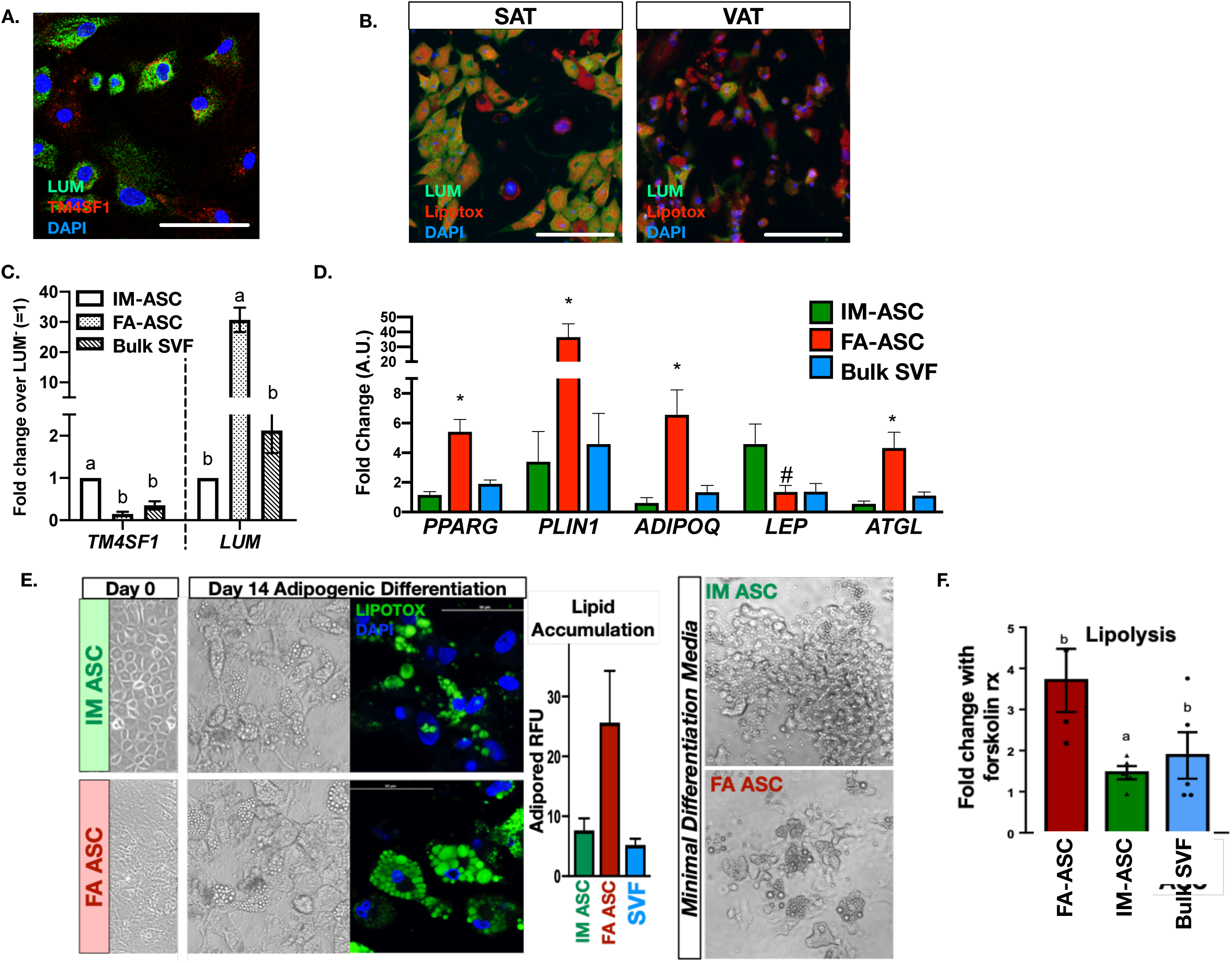
Distinct metabolic profiles of *in vitro* differentiated IM-ASC and FA-ASC. (A) Immunofluorescence evaluation of bulk VAT ASC cultures showing differential expression of LUM (FA- ASC) and TM4SF1 (IM-ASC) in distinct cell ASC types. (B) Immunofluorescence evaluation of bulk SAT and VAT ASC cultures differentiated into adipocytes and stained with LUM (FA-ASC; green) and LipidTox to identify differentiated, lipid-filled adipocytes. (C) Gene expression analysis by qRT-PCR of FACS sorted VAT IM-ASC and FA-ASC in comparison to bulk SVF of mixed ASC from VAT for IM- ASC (*TM4SF1*) and FA-ASC (*LUM*) specific genes. Fold changes were calculated relative to IM-ASC. (D) Gene expression analysis by qRT-PCR of differentiated IM-ASC, FA-ASC, and bulk VAT ASC cultures of key adipogenic genes. Fold changes were calculated relative to IM-ASC. (E) Adipogenesis of FACS sorted VAT IM-ASC and FA-ASC with quantitation of lipid accumulation by LipidTox and Adipored staining (n=4 subjects). Right: bright field images of adipogenic differentiation in the absence of insulin and TZD. (F) Lipolysis assays of differentiated IM-ASC, FA-ASC, and bulk VAT ASC cultures. Fold changes were calculated using basal, non-stimulated cells.

*In vitro* expanded and adipogenically differentiated ASC are a widely used tool for modeling adipocytes and a potential source of cells for cell-based therapy for metabolic disease in humans. To compare the phenotypes of ASC derived from fresh AT SVF with *in vitro* expanded ASC, flow cytometry was performed on paired VAT and SAT bulk ASC cultures from obese subjects. Enrichment of CD31^-^ TM4SF1^+^ IM-ASC was seen VAT, but not SAT, for all subjects (Figure 5A-B). Small populations of CD31^+^TM4SF1^+^ EC were observed in some cultures consistent with our IF in whole AT. Consistent with overlap between IM-ASC and CD9^hi^ ASC, flow cytometry showed higher CD9 expression in TM4SF1^+^ IM-ASC compared to TM4SF1^-^ FA-ASC (Figure 5C). This demonstrates stable maintenance of depot specific IM-ASC and FA-ASC subpopulations with *in vitro* expansion of VAT and SAT ASC.

**Figure 5.**
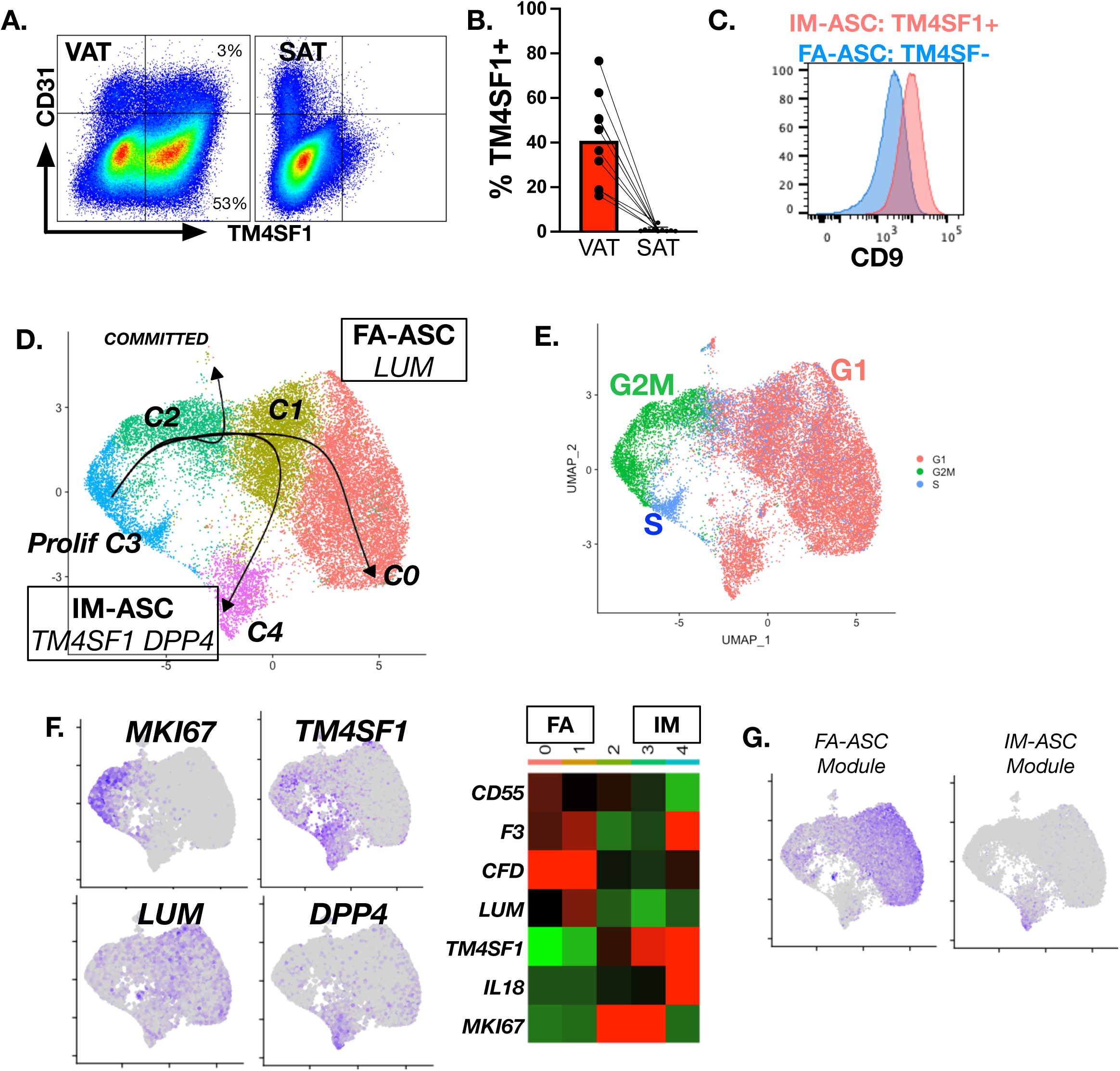
Retention of depot specific ASC diversity in *in vitro* cultures. (A) Flow cytometry identification of VAT specific CD31^-^TM4SF1^+^ (IM-ASC) and CD31^-^TM4SF1^-^ (FA-ASC) in *in vitro* expanded SVF cultures from obese patients. (B) Quantitation of TM4SF1 in paired VAT and SAT *in vitro* SVF cultures from subjects with obesity (n=8). (C) Expression of CD9 in CD31^-^TM4SF1^+^ (IM-ASC) and CD31^-^TM4SF1^-^ (FA-ASC). (D) Single-cell RNAseq (scRNAseq) of *in vitro* cultured bulk VAT ASC from subjects with obesity (n=4). Trajectory analysis performed with Celltrace with origin at proliferating C3. (E) Cell cycle analysis of scRNAseq from *in vitro* cultured VAT ASC. (F) Feature plots and heatmaps for proliferative (*MKI67*), IM-ASC (*TM4SF1, DPP4*), and FA-ASC (*LUM*) specific genes derived from scRNAseq of i*n vitro* cultured VAT ASC (G) Feature plots of expression levels for FA-ASC and IM-ASC gene modules in cultured ASC scRNAseq dataset.

To complement this analysis, we performed scRNAseq on bulk VAT ASC cultured 3-5 passages (Figure 5D, Table S4). Cell cycle analysis supports a common LUM^-^ DPP4^-^ proliferating stem cell (C3) in S-Phase followed by G2M cells (C2) and several types of G1 phase cells (C0, C1, C4) (Figure 5E). Consistent with this C2 and C3 had high expression of proliferation markers (*MKI67*) cell population and were LUM^-^ DPP4^-^TM4SF1^low^. Among the G1 phase cells, distinct LUM^-^TM4SF1^high^DPP4^+^ ASC (C4; IM-ASC), TM4SF1^low^LUM^+^ ASC (C0,C1; FA-ASC), and a small population of committed progenitors expressing early differentiation genes (*PPARG, FABP4*) were identified (Figure 5F). Computing scores for IM- and FA-ASC gene modules confirmed analogous populations of LUM^+^ FA-ASC (C0, C1) and TM4SF1^+^ IL18^+^ IM-ASC (C4) in the *in vitro* cultures (Figure 5G). Trajectory analysis (Slingshot ^33^) identified differentiation pathways that generated both LUM^-^TM4SF1^high^DPP4^+^ ASC (C4; IM-ASC), LUM^+^ ASC (C0,C1; FA-ASC) (Figure 5D). Overall, our *in vitro* data concurs with *in vivo* data demonstrating distinct lineages of that are stable with *in vitro* culture and expansion.

### Mature adipocyte heterogeneity consistent with ASC diversity

Heterogeneity of human adipocytes has been reported by others and is implied by our results by the adipogenic differentiation of distinct ASC subtypes *in vitro.* We examined our snRNAseq dataset for evidence of IM- and FA-like adipocytes *in vivo*. Analysis of adipocytes (*ADIPOQ, PLIN1, G0S2*) found enrichment of IM-ASC genes (*TM4SF1*, *IL18*) in VAT, but not SAT adipocytes, suggesting that IM-ASC differentiate into adipocytes (Figure 6A).

**Figure 6.**
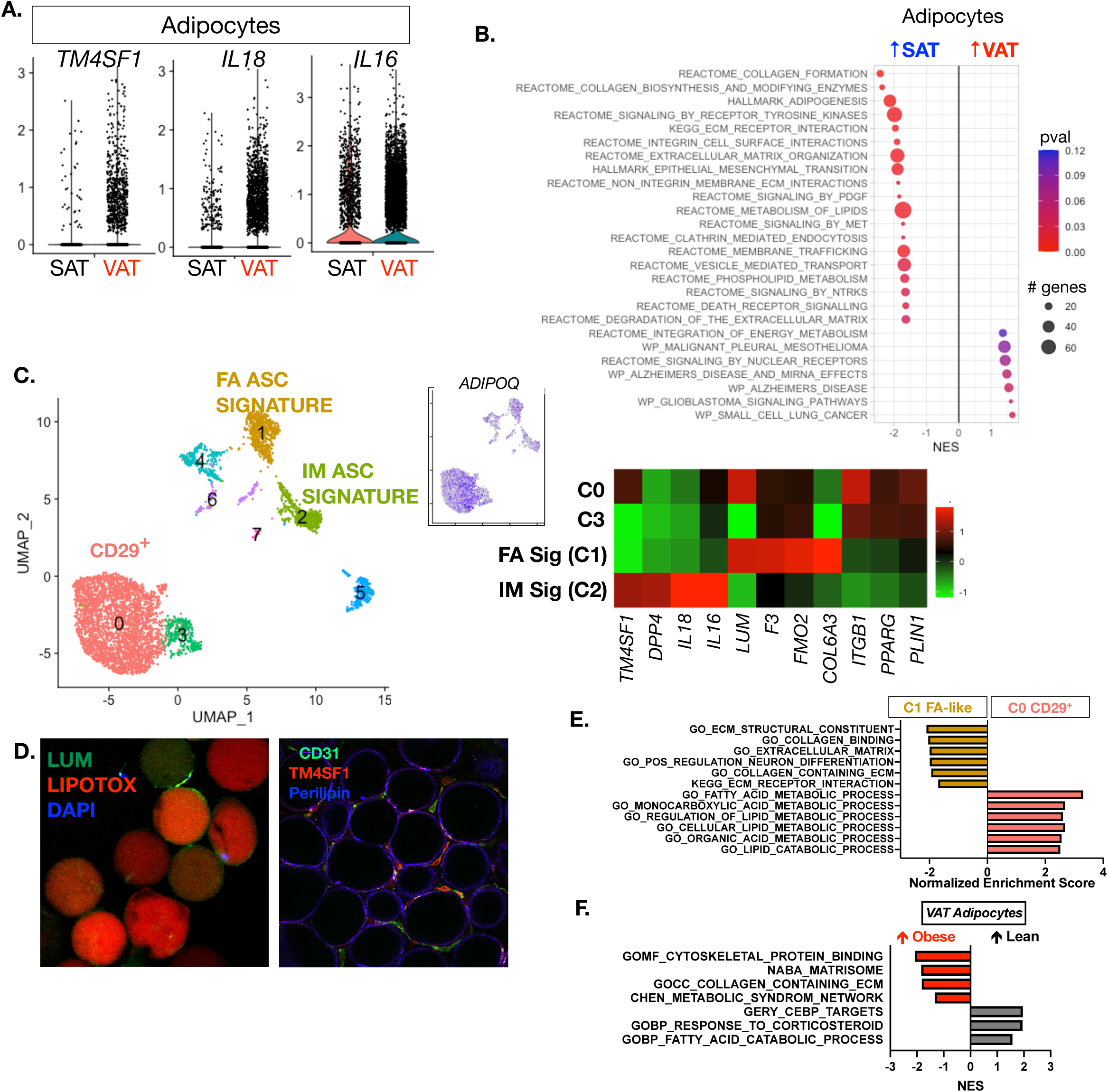
Adipocyte diversity in human SAT and VAT. (A) Depot specific differences in expression of *TM4SF1, IL16,* and *IL18* in adipocyte (ADIPO) cluster in snRNAseq dataset. (B) Pathway analysis (GSEA) of DE genes between SAT and VAT adipocytes (ADIPO). (C) Subanalysis of adipocytes (expressing *ADIPOQ*) in core dataset and heatmap of key DE genes. (D) Left: Immunofluorescence identification of LUM^+^ adipocytes (FA-ASC related) isolated after collagenase digestion of VAT followed by confocal microscopy imaging. Intracellular lipids were stained with LipidTox (red). Right: Immunofluorescence identification of TM4SF1^+^ adipocytes in whole mount adipocytes by confocal microscopy (*right*). Perilipin (blue) was used to identify adipocytes and CD31 to identify endothelial cells. (E) Pathway analysis (GSEA) comparing DE genes between C1 FA-ASC like and C0 CD29^+^ adipocyte clusters. (F) GSEA pathway analysis comparing DE genes between obese and lean subjects in VAT adipocytes.

Pathway analysis comparing DE genes between SAT and VAT adipocytes demonstrated enrichment for ECM, integrin, and adipogenic gene pathways in SAT adipocytes and enrichment for mesothelial and nuclear receptor genes in VAT adipocytes (Figure 6B). Subset analysis of the mature adipocyte population revealed 7 distinct mature adipocyte subpopulations within the higher-level mature adipocyte cluster (Figure 6C, Table S5). The largest adipocyte type (C0) expresses *ITGB1/CD29,* but not *LUM* or ASC markers. Analysis of expression of LUM^+^ ASC markers identified a cluster of cells (C1) with high expression of multiple markers of LUM^+^ FA-ASC progenitors (*DCN, F3, LUM, NEGR1*).

Another adipocyte cluster (C2) expressed markers of IM-ASC (*TM4SF1, DPP4*). The observation of distinct adipocyte types that retain expression of markers of specific ASC is consistent with our *in vitro* differentiation observations. Consistent with this, IF assessment of collagenase-isolated primary VAT adipocytes identified LUM^+^ and LUM^-^ adipocytes (Figure 6D). In addition, TM4SF1^+^ adipocytes were identified in VAT consistent with an origin from IM-ASC. GSEA comparing FA-ASC-like adipocytes with CD29^+^ adipocytes showed an increase in ECM pathways in FA-ASC adipocytes (Figure 6E). Our dataset permits us to examine obesity dependent genes in VAT, but not SAT adipocytes. Pathway analysis of DE genes identified an increased in matrisome, collagen and genes in the metabolic syndrome network that are increased in obese VAT adipocytes (Figure 6F). In contrast, lean VAT adipocytes are enriched for fatty acid catabolic genes and genes related to corticosteroid responses.

### AT endothelial (EC) and pericyte (PC) diversity

Previous scRNAseq experiments underrepresent the number of EC in AT and we were struck by the identification of multiple EC subtypes in our scRNAseq data defined primarily by expression of *VWF*. Sub-analysis of EC and PC populations was performed to identify distinct subpopulations (Figure 7A, Table S6). Significant expression of genes involved in lipid metabolism was observed in all EC and PC subtypes including *PPARG, FABP4,* and *ZNF423* (Figure 7B-C). These observations are consistent with prior data suggesting that PC or EC may be adipogenic precursors. ^20, 34, 35^

**Figure 7.**
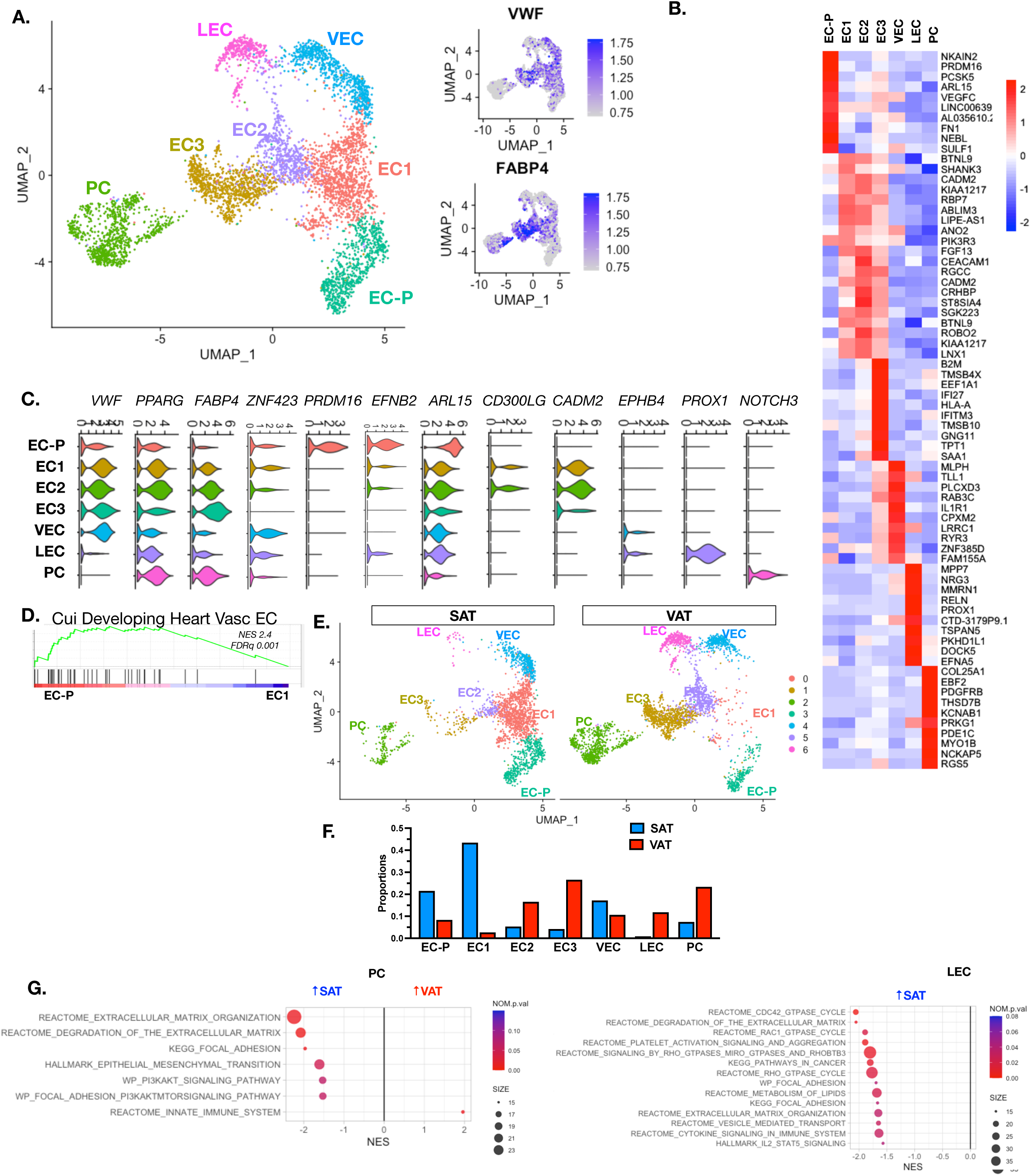
Diversity of endothelial and pericyte populations in AT. (A) UMAP plot of subanalysis of EC, EC-P, and PC populations in core snRNAseq dataset with feature plots for *VWF* and *FABP4*. (B) Heatmap of hallmark genes for EC subpopulations in 7A. (C) Violin plots for hallmark genes for EC subpopulations in 7A. (D) Pathway analysis (GSEA) of DE genes between EC-P and EC1 cells. (E) UMAP plots of EC/PC split by depot. (F) Proportions of EC/PC subtypes in SAT and VAT. (G) Pathway analysis (GSEA) of DE genes between SAT and VAT in PC (*left*) and LEC (*right*).

A putative adipose EC progenitor cell (EC-P) was identified with markers of an immature phenotype based on low *VWF* expression, expression of early arterial EC markers (*EFNB2*)^36^, and *NOTCH* signaling genes (*NOTCH4, DLL4, JAG2, HEY1*) critical for EC development. Pathway analysis comparing EC-P with mature EC populations (EC1) showed a significant enrichment in genes involved in vascular development in EC-P (Figure 7D, Table S7). Notably, the strong DM risk gene *ARL15* was highly enriched in EC-P. Surprisingly *PRDM16* which is involved in generating arterial EC in addition to its role in brown adipogenesis^37^ was specifically expressed in EC-P. EC1 and EC2 manifested high VWF expression as well as expression of CDH13/T-Cadherin – an adiponectin receptor. Venous EC (VEC) were identified with high expression of VEC markers *NR2F2*, *EPHB4*, and *SNTB2*.^37^ A *lymphatic* EC (LEC) was identified with high *PROX1* and no *VWF* expression.^38, 39^ NOTCH3^+^ pericytes (PC) were identified. Expression of *CD146*/*MCAM,* a marker commonly used to identify PC ^40, 41^, was not exclusive to PC - challenging the sole use of CD146 as a PC marker.

The distribution of PC and EC cell types were compared between VAT and SAT (Figure 7E-F). The proportions of EC-P and EC1 were significantly increased in SAT compared to VAT consistent with reports of increase angiogenic capacity in SAT relative to VAT. EC2, EC3, LEC, and PC were increased in VAT compared to SAT. Comparison of PC gene expression between VAT and SAT showed an enrichment of pathways involved in ECM organization, degradation, and epithelial mesenchymal transition in SAT PC and an enrichment of innate immune system genes in VAT PC (Figure 7G). Similar enrichment in ECM pathways, focal adhesion pathways, and lipid metabolism were observed in SAT LEC compared to VAT.

### Immune cell profiles in VAT and SAT

Sub-analysis of the immune cell populations identified macrophage (MAC; *MERTK, CD163*), dendritic cell (DC; *CD1D, FLT3*), T cell, and NK cell (*SKAP1*) populations (Figure 8A, Table S7).

**Figure 8.**
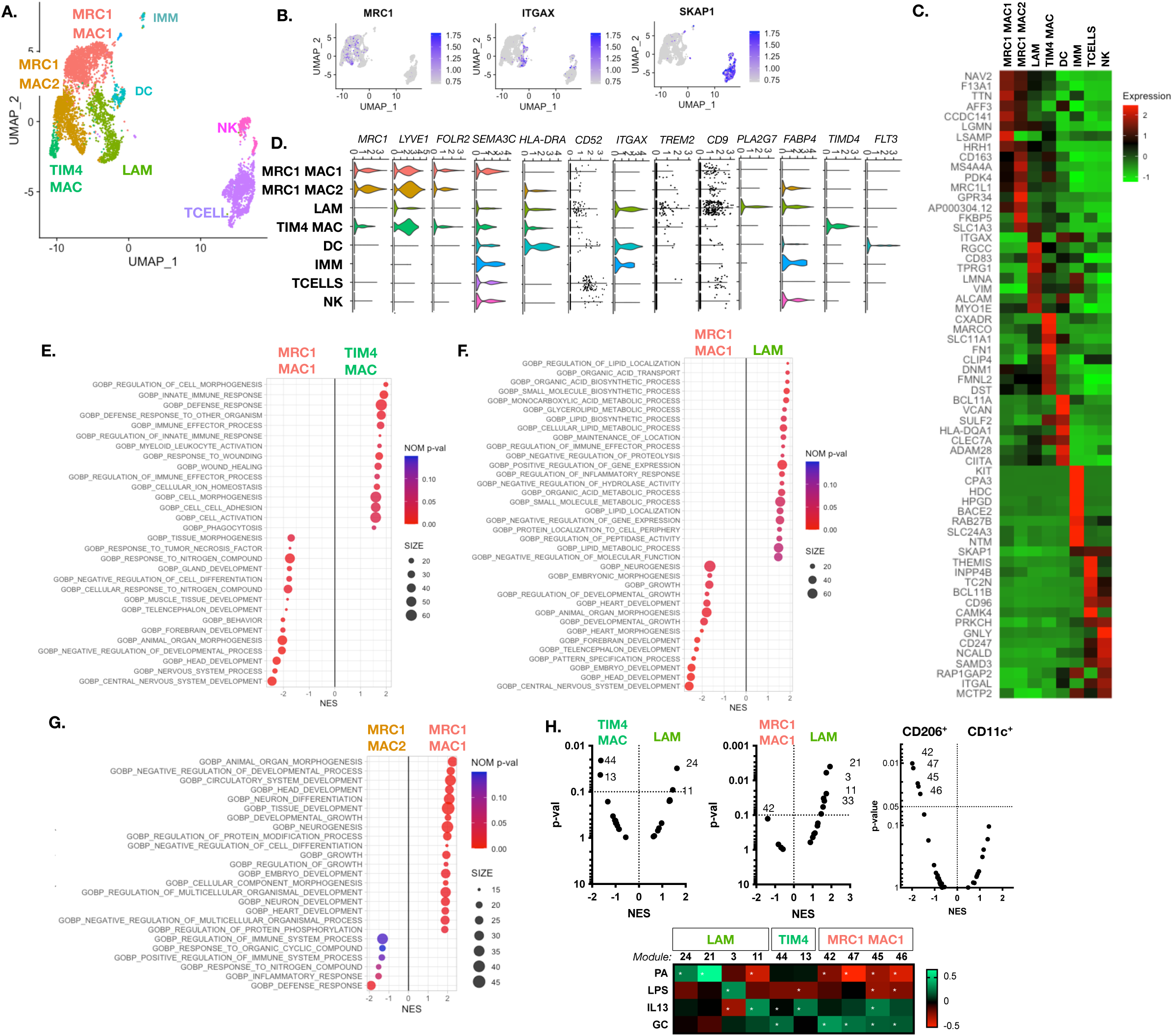
Immune cell diversity in SAT and VAT. (A) Subset analysis of immune cells in core snRNAseq dataset. Two subsets of *MRC1* macrophages identified along with *TIM4* and Lipid Activated Macrophages (LAM). (B) Feature plots of hallmark genes of *MRC1* (CD206) macrophages, *ITGAX* (CD11c) macrophages, and *SKAP1* T cells. (C) Heatmap of hallmark genes for immune cell populations. (D) Violin plots of key macrophage (MAC) and dendritic cell (DC) markers. (E) Pathway analysis (GSEA) of DE genes between MRC1 MAC1 and TIM4 MAC. (F) Pathway analysis (GSEA) of DE genes between MRC1 MAC1 and LAM. (G) Pathway analysis (GSEA) of DE genes between MRC1 MAC1 and MRC1 MAC2. (H) GSEA analysis of AT macrophages based on human macrophage activation gene modules. Module activating conditions noted for palmitic acid (PA), LOS, IL13, and glucocorticoids (GC) from ^48^. * indicates significant induction of gene module by the activating condition.

Focusing on the macrophage populations, four main macrophage subtypes were identified. Aligning with previous studies ^42^, a lipid associated macrophage (LAM) subtype was identified with high expression of *ITGAX, TREM2, CD9,* and *CD52* (Figure 8B-D). Platelet activating factor acetylhydrolase (PLA2G7) which has been recently shown to be decreased in AT in humans under calorie restriction is specifically expressed in LAM macrophages.^43^ *MRC1/CD206* were identified as enriched in two subpopulations (MRC1 MAC1, and MRC1 MAC2) that also express resident macrophage markers *F13A1, PDGFC,* and *LYVE1* aligning with our previous flow-based assessments ^44^ and other studies^45^. A *TIMD4* macrophage population was observed with lower *MRC1* expression consistent with prior reports of a unique regulatory TIMD4^+^ tissue resident macrophages ^46, 47^ .

Comparison of DE genes TIM4 MAC and MRC1 MAC1 identified enrichment of pathways for innate immune responses and myeloid activation in TIM4 MAC while pathways involved in tissue morphogenesis and development were enriched in MRC1 MAC1 (Figure 8E). Comparison of DE genes LAM and MRC1 MAC1 identified enrichment of pathways for lipid metabolicm and inflammatory responses in LAM and the enrichment of genes involved in tissue development in MRC1 MAC1 (Figure 8F). Comparison of DE genes between MRC1 MAC1 and MRC1 MAC2 cells showed an enrichment for genes related to development and neurogenesis in MAC1 and enrichment for pathways involved in immune responses in MAC2 (Figure 8G).

A spectrum of human macrophage activation has been described and gene modules that respond to a range of stimuli have been identified.^48^ GSEA analysis was used to identify which gene modules representing the spectrum of *in vitro* macrophage activation were enriched in the different ATM subpopulations (Figure 8H). Comparison of LAM with TIM4 MAC demonstrated the induction of lipid dependent gene modules in LAM (Modules 24, 11) while TIM4 MAC had enrichment of gene modules that are induced by glucocorticoids (Module 44) and Th2 cytokines IL13 and IL4 (Module 13). Comparison of LAM with MRC1 MAC1 identified enrichment in for gene modules (Modules 21, 3, 11, 33) that are up and down regulated by fatty acids (palmitic acid, oleic acid, stearic acid) while MRC1 MAC1 cells are enriched for module 42 which is induced by glucocorticoids. The enrichment of glucocorticoid modules was recapitulated in FACS sorted MRC1/CD206^+^ human ATMs (Modules 42, 47, 45, 46) relative to ITGAX/CD11c^+^ ATMs from our recently published study.^44^ This demonstrates that canonical M1 and M2 modules do not differentiate the different ATM types and that enrichment of glucocorticoid induced genes is a feature of MRC1/CD206^+^ human ATMs.

## Discussion

This study provides a novel resource for interrogation of AT cell types using single nuclear RNAseq approaches from human subjects. Our datasets align with and complements similar datasets from humans and mice that expand our understanding of diverse AT cell types and states. ^7, 11, 49^. We have focused on depot-specific differences between SAT and VAT cell types– two critical depots with different functions and associations with metabolic dysfunction in obesity. Our main findings are the presence of diverse subtypes of endothelial cells in human AT with features of lymphatic, venous, and arterial EC, identification of a PRDM16 expressing EC population with features of an EC progenitor, the identification of a VAT-specific ASC type (IM-ASC) with mesothelial-like features, and the demonstration and validation that multiple ASC types can give rise to adipocytes that retain features of their progenitors. Comparison with our dataset and scRNAseq datasets (e.g. Vijay et al^1^) emphasize multiple cell types underrepresented in scRNAseq datasets beyond adipocytes.

Comparison of our ASC data to other published data highlight important similarities and differences in approach that will need to further clarified experimentally especially from human samples. Based on transcriptomics analysis of flow-sorted murine ASC, Marcelin et al. described two PDGRFA+ subpopulations distinguished by high or low expression of CD9.^18^ CD9^hi^ cells expressed high levels of ECM molecules, while CD9^lo^ cells expressed high levels of adipogenic genes and manifested an adipogenic phenotype upon differentiation in vitro. Furthermore, CD9^lo^ cells shifted from an adipogenic to a fibrotic phenotype and were decreased in frequency in response to HFD. Marcelin et al. corroborated these findings in humans, finding decreased CD9^lo^ cells (analogous to our FA-ASC) and increased CD9^hi^ cells (analogous to our IM-ASC) in insulin-resistant humans with obesity, frequencies that correlated with the degree of VAT fibrosis. Notably, our data support a similar shift in FA-ASC phenotype from adipogenic to fibrotic phenotype in obesity, consistent with Marcelin et al.’s observations of CD9^lo^ ASC in mice in response to HFD.

Merrick et al. used scRNASeq to describe seven ASC-like populations in mice, three of which expressed canonical ASC markers *PDGFR1, Sca1, CD34*, and *Thy1* (DPP4^+^, ICAM1^+^, CD142^+^).^2^ Two of these subpopulations correlated with similar ASC populations in human SAT (DPP4^+^, ICAM1^+^). Merrick described DPP4^+^ ASC as a multipotent progenitor that gave rise to a committed adipogenic ICAM1^+^ progenitor. CD142^+^ cells also manifested adipogenic potential, but demonstrated transcriptional similarities to anti-adipogenic “A-regs” described by Schwalie et al..^3^ DPP4+ cells were noted to be decreased, and CD142^+^ cells increased, in obese compared to lean mice. Merrick et al.’s study reports depot-specific differences in these populations, with CD142^+^ cells increased in eWAT relative to iWAT in mice, consistent with similar observations by Schwalie et al. In contrast, Merrick et al. decribe that mouse and human CD142^+^ had high adipogenic potential, while Schwalie et la. found these cells have low adipogenic potential. The authors concluded that this discordance was the result of differences in FACS isolation strategies. Our IM-ASC overlap transcriptionally with DPP4^+^ ASC, identified by Merick’s, except that they found these cells in human SAT (not VAT) and described a decrease in frequency in HFD in eWAT, which diverges from our observations in humans. Merrick et al. also localized DPP4+ cells to the reticular interstitium, suggesting similarities to a mesothelial sub-anatomic localization that we observed for at least a portion of IM-ASC. Our IM ASC also resemble mesothelial-like cells by Hepler et al., and MSLN^+^ VP1/VP3 of Vijay et al.^1^, reinforcing a mesothelial origin for some of our IM-ASC. Our observations of VAT-specificity for IM-ASC agree with Vijay et al. for VP1/VP3 shown to have high mitochondrial activity. Our FA-ASC are analogous to CD142^+^ ASC described by Merrick et al. (except that Merrick found more CD142 in eWAT than iWAT), and CFD^+^ SP1/SP4/VP4 populations of Vijay et al. that correlate with type 2 diabetes.

In a depot-specific characterization of adipocyte progenitor cell (APC) diversity in human AT, Raajendiran *et al.* used scRNAseq to identify three distinct Lin- CD29+ APC subpopulations in the omentum (VAT), the abdominal subcutaneous (ASAT), and the subcutaneous gluteofemoral (GFAT) AT from subjects with obesity. APC subpopulations were flow-sorted from SFV for further metabolic interrogation *in vitro* based on their level of CD34 expression: high (CD34^hi^), low (CD34^lo^), and negative (CD34^-^) expression. Transcriptionally, CD34^hi^ APCs highly overlap with our FA-ASC (*LUM, DCN, APOD*, *FBLN1*, and collagens), while CD34^lo^ gene profile (*TM4SF1, MSLN, ITLN1, PLA2G2A, UPK3B*) coincides with our IM-ASC. In contrast to our study, CD34^hi^ were more abundant in VAT compared to ASAT, while CD34^lo^ APCs were more abundant in GFAT compared with VAT (% of Lin- cells) and were not VAT-specific as our IM-ASC. Similar to our study, Raajendiran *et al.* evaluated the adipogenic capacity and metabolic phenotype of the distinct APCs subpopulations *in vitro.* All three APC subpopulations underwent adipogenesis and lipid loading, but there was no difference in overall lipid accumulation between the APC subtypes in VAT, ASAT and GFAT; however, CD34 ^hi^ adipocytes had greater fatty acid uptake compared with CD34 ^lo^ and CD34^-^ in all depots. In contrast, we did observe increased lipid accumulation in *in vitro* differentiated FA-ASC compared with IM-ASC. CD34^hi^ had increased lipolysis compared with CD34lo and CD34-, and higher gene expression of *ATGL* and *LIPE*, similar to our findings with FA-ASC adipocytes. Notably, Raajendiran *et al.* report and increased proportion of CD34^hi^ APCs in subjects with DM, which implies an interaction between the transcriptional and metabolic phenotype of distinct APCs with AT dysfunction observed in DM.

Much of the discordance between murine and human ASC subpopulations may arise from species differences in expression of canonical ASC markers. Supporting a role for an adipogenic mesothelial cell, Hepler et al. ^5^ described in mice a population of “mesothelial-like cells” (“Cluster 3”, “FIP”) defined by expression of MSLN and other mesothelial markers that co-expressed PDGRFb and other ASC markers. A number of studies use PDGFRa and PDGFRb as markers for ASC. These markers are relatively restricted to ASC in mice, but in humans are promiscuous, and not uniformly expressed by all ASC. Emont et al. demonstrate PDGFRa and PDGFRb expression in humans not only by ASC, but also by mesothelial cells, mature adipocytes, T-cells, macrophages, and endothelial cells.^7^ These observations suggest that extrapolation of murine studies of ASC defined by these markers may not apply to humans.

Other studies have shown differences in sn and sc RNAseq data that may contribute differences in our dataset and others.^6, 50^ Indeed the higher degree of concordance of our data with that of Emont et al., who performed snRNASeq, compared with Vijay et al., who used scRNASeq, support such differences. In addition, while some studies such as Vijay *et al.* suggest a larger number of human ASC subtypes than our current data, few of their computationally defined types have been experimentally or functionally validated. The number of patients studied and the depth of sequencing are important considerations in sc/snRNAseq. We observed that interpatient variability was a significant factor in resolving cell subpopulations from snRNAseq data, with a greater effect than disease (DM status) or obesity status. These observations suggest that very large patient numbers will be necessary to adequately power snRNAseq studies to define cell populations and gene expression patterns specific for clinical characteristics such as disease, obesity status, age, gender, and others. We also observed marked differences in cell subpopulations frequencies between VAT and SAT, suggesting qualitatively different biology of these AT depots.

The role of mesothelial cells in AT is controversial. Published data support adipocyte progenitor cells of mesothelial origin.^51,52,53^, with WT1^+^ mesothelial like cells serving as adipocyte progenitors.^54^ Rosen’s group recently demonstrated that KRT19 may represent a more specific mesothelial marker than WT1 however, and that murine KRT19^+^ cells are not adipogenic.^55^ In a separate report, Emont et al corroborated in humans the presence of a MSLN^+^KRT19^+^PDGFRA^-^ mesothelial cell population, but did not provide functional data addressing adipogenic potential of these human cells. Indeed, species-specific differences in a mesothelial contribution to adipocyte ontogeny may exist. In support of such differences, Emont et al. did not find significant differences in the frequencies of ASC subpopulations between depots in mice, in contrast to humans, in which specific subpopulations were markedly enriched in VAT (hAd2, hAd6) or SAT (hAd1, hAd3, hAd4, hAd7). Notably, hAd 3 and 4 enriched in SAT expressed high levels of genes associated with ‘triglyceride biosynthesis’ and fatty acid desaturases respectively, consistent with an adipogenic phenotype for human SAT adipocyte progenitors. In addition to species-specific differences, it is possible that our IM-ASC may consist of both mesothelial and non-mesothelial cells. Our data demonstrate that KRT19 is expressed at high frequency by IM-ASC, but at lower frequencies by FA-ASC and T-cells. In addition, we identify KRT19^+^ and KRT19^-^ IM-ASC, suggesting that IM-ASC may consist of both mesothelial and non-mesothelial-like cell subpopulations, explaining our observations of anatomic localization of these cells in both the mesothelial lining and parenchyma of AT, and providing opportunity for finer resolution of our IM-ASC population.

The question of whether ASC cells maintain subpopulation phenotypes with in vitro culture and passaging has important implications for investigative and translational applications. Our observation that depot-specific IM- and FA-ASC subpopulation frequencies and transcriptional phenotypes are maintained in culture are consistent with data from Emont et al. demonstrating similar maintenance of ASC subpopulations in culture. Similar to Emont et al., our data also support that subpopulations of mature adipocytes manifest transcriptional phenotypes similar to ASC subpopulations, suggesting that these cells derive from corresponding ASC subpopulations.

Our evaluation of AT immune cells demonstrates a range of macrophages that align with recent surveys of mouse and human AT leukocytes.^47^ We note that our dataset did not identify significant numbers of leukocytes such as mast cells which have been previously identified in human AT^56^ or eosinophils which are prominent in murine AT but have scant evidence for significant numbers in human AT. ^57^

Our dataset is currently underpowered to identify the full complement of obesity and diabetes specific gene expression differences on a cell type specific basis. Integration with current and future datasets may add to the power to discriminate gene expression changes that delineate cell type specific changes in gene expression associated with disease. Future steps will also include the addition of single nuclear ATAC strategies and multiomic analysis to identify the regulatory regions that are responsible for the gene expression changes as has been recently performed for skeletal muscle in diabetes.^58^

## Materials and Methods

### Human subjects

Human subjects were enrolled with Institutional Review Board approval at University of Michigan and Ann Arbor Veterans Affairs Healthcare System. Informed consent was obtained and human subjects research carried out in accordance with The Code of Ethics of the World Medical Association, the Belmont Report, and the Declaration of Helsinki. VAT from the greater omentum and SAT from the anterior abdominal wall were collected at the beginning of operation from obese subjects undergoing bariatric surgery and from lean subjects undergoing general surgery procedures (Table 1). Subjects with type 2 diabetes (DM) were defined by clinical diagnosis requiring medication and hemoglobin A1c (HbA1c)>=6.5%. Non-diabetic (NDM) subjects were defined by no clinical history of diabetes and HbA1c<5.7% per American Diabetes Association criteria.

### AT nuclei isolation and single-nuclei RNA sequencing

Nuclei isolation from human VAT and SAT was adapted from a previous protocol.^19^ Briefly, 400-500mg samples of frozen human VAT or SAT were washed with sterile RNAse-free cold 1X DPBS and minced with a scalpel in a petri dish on ice. Samples were homogenized in a 15 mL precooled glass dounce homogenizer (10 strokes with pestel A followed by 15 strokes with pestel B) with 1.5mL of homogenization buffer [5mM MgCl2, 10mM Tris Buffer (pH 8.0), 25mM KCl, 250mM sucrose, 1X protease inhibitor, 1uM DTT, 0.4 U/uL Ribolock Rnase Inhibitor (40U/μL), 0.2U/uL Superasin (20U/uL), 0.1% triton X-100] and strained with pre-wet 100 µm and 40 µm filters into a 50mL conical tube. Next, each sample was transferred to two 1.5 mL pre-chilled microcentrifuge Rnase-free tubes and centrifuged at 500rcf, 4°C, 5 minutes. Supernatant was removed with a pipette, leaving approximately 50uL remaining, which was resuspended in 500uL of 1% BSA-DPBS including 0.2U/μl of Rnase Inhibitors. Before proceeding with fluorescence activated cell sorting (FACS), a subsample of nuclei was used to assess the overall quality of the nuclei by staining with trypan blue and visualized by light microscopy. Nuclei were stained with propidium iodide (PI) for sorting (Invitrogen, Cat#V13241, 10ug/mL) at a 1:100 dilution, leaving approximately 20-30uL of sample as unstained control. Each sample was transferred into a pre-coated 5 mL polystyrene flow cytometry tube and sorted using a 100uM nozzle in a FACSAria III Cell Sorter (BD Biosciences Inc., Franklin Lakes, NJ USA). Sorting strategy included doublet discrimination and selection of intact nuclei by subgating on PI^+^ nuclei. PI^+^ nuclei was sorted directly into a 1.5 mL microcentrifuge tube containing 20uL of 1% BSA-DPBS and 0.2U/μl of Rnase Inhibitors. After nuclei isolation, samples were centrifuged at 100rcf, 4°C, 6 minutes, supernatant was pipetted off leaving approximately 50uL of resuspended nuclei. Single-nuclei suspensions were subjected to final nuclei counting on Countess II Automated Cell Counter (ThermoFisher Scientific Inc., Waltham, MA USA) and diluted to 700 -1000 nuclei/ul. 3’ single nuclei libraries were generated using the 10X Genomics Chromium Controller and following the manufacturer’s protocol for 3’ V3.1 chemistry with NextGEM Chip G reagents (10X Genomics Inc., Pleasanton, CA, USA). Final library quality was assessed using the Tapestation 4200 (Agilent Inc., Santa Clara, CA USA) and libraries were quantified by Kapa qRTPCR (Roche Sequencing and Life Science Inc., Wilmington, MA, USA).

### Single-cell suspension preparation from human VAT ‘bulk’ preadipocytes

Preadipocytes derived from VAT-SVF of four DM patients were cultured in vitro until confluency in a T175 flask using preadipocyte medium. Single-cell suspension was obtained by detaching preadipocytes with 5 mL of 0.25% trypsin (ThermoFisher Scientific Inc./Gibco) for 3 min at 37C, followed by centrifugation (500 rcf, 5 min, 4C), and resuspension with 1mL of 1X DPBS with 0.04% BSA with gentle mixing using a wide-bore tip. A pre-wet 40 um cell strainer was used to remove any remaining cell debris and large clumps. A 10 uL sub-sample was obtained to determine viability and the cell concentration using trypan blue and automated cell counter. All samples had cell viability >95%. Cell suspensions were centrifuged at 300 rcf for 5 min, supernatant removed, and single-cell suspensions diluted to 1000 cells/uL using 1X DPBS with 0.04% BSA. Single-cell suspensions were gently mixed 10-15 times until cells were completely resuspended and placed on ice. Samples were immediately transported to the Advanced Genomics Core Facility at University of Michigan for proceeding with the 10x Genomics single-cell protocol.

### Single nuclear and cell RNA sequencing data analysis pipeline

Pooled libraries were subjected to 150 bp paired-end sequencing according to the manufacturer’s protocol (NovaSeq 6000, Illumina, Inc., San Diego, CA, USA). Bcl2fastq2 Conversion Software (Illumina Inc.) was used to generate de-multiplexed Fastq files and CellRanger Pipeline (10X Genomics Inc.) was used to align reads and generate count matrices using whole genome reference libraries. Ribosomal and mitochondrial genes were removed prior to analysis with Seurat V4.0. Nuclei were filtered to remove low and high UMI cell/nuclei and background subtracted with SoupX. Integrated analyses were performed with Seurat sctransform algorithms. DE genes identified with MAST2. Pathway analysis of DE genes performed using GO terms, Hallmark terms, and KEGG pathways in MsigDB. Dimensional reduction with UMAP. Module scores for the top 100 enriched genes for different published cell types were computed for each cell type and presented to identify enrichment of different cell profiles compared to published data sets. See Supplemental Tables for gene expression lists. Single nuclear RNAseq data deposited in the Single Cell Portal at Accession <u>SCP1903</u>. Single cell RNAseq data from cultured human VAT preadipocytes deposited at Accession <u>SCP1905</u>.

### Human SVF isolation, FACS strategy

Human VAT and SAT biopsies (30-50g) were minced and digested with 3mg/mL type II collagenase solution (ThermoFisher Scientific Inc./Gibco) in HBSS containing calcium chloride, magnesium chloride, and glucose (ThermoFisher Scientific Inc., Cat#14025092) and 0.5% fatty acid free- BSA (Sigma Aldrich Inc., Burlington, MA USA Cat# A8806) in a shaker at 37°C, 30 minutes. Digests were filtered through 100uM strainers (Corning Inc./Falcon, Corning, NY USA), rinsed with HBSS (Ca^++^-free, Mg^++^- free) supplemented with 0.5% BSA, centrifuged at 300rcf, at 4 °C, 10 minutes, and SVF pellet resuspended in 1X red blood cell lysis buffer (RBC; 155mM NH4Cl, 10mM KHCO3, 1mM EDTA), 25 °C, 6 min. After washing with cold 1X DPBS, samples were centrifuged, resuspended in 0.5% BSA DPBS (FACS buffer) and counted manually using trypan blue staining (0.4%, Gibco Inc., Cat# 15250-061).

Cells were immunolabeled with CD45-PE (Biolegend Inc., San Diego, CA USA, Cat# 304008), CD31- PE/Cy7 (Biologend Inc., Cat# 303117) and TM4SF1-APC (Miltenyi Biotec Inc., Bergisch Gladbach, Germany, Cat# 130-112-905) at 8°C, 30 minutes in FACS buffer. Cells were washed twice with 3mL of FACS buffer, stained with DAPI (1ug/mL) 25°C, 10 minutes, washed once with FACS buffer, and strained in a pre-wetted CellTrics 50um sterile filter (Sysmex Inc. Kobe, Japan). FACS controls included unstained cells and single stain samples. FACS was performed in a Synergy SY3200 cell sorter at the University of Michigan Flow Cytometry Core. We used doublet discrimination strategies by eliminating small particles with an area plot of forward scatter (FSC) versus side scatter (SSC), followed by height versus FSC/SSC area. Subsequent plots gated based on cell viability (DAPI versus SSC area) and double negative CD45 and CD31 cells (non-immune, non-endothelial cells). A subsample of non-sorted SVF from each patient was used as control in subsequent analyses. Flow cytometry analysis was performed with FlowJo. After FACS, cells were re-counted and washed 1X with FACS buffer. TM4SF1 surface marker was used to discriminate distinct ASC subpopulations (CD45^-^CD31^-^TM4SF1^+^ and CD45^-^CD31^-^TM4SF1^-^).

### ASC culture and differentiation

Bulk ASC were isolated from VAT stromal vascular cell fraction (SVF) of obese DM and NDM patients, as described ^12, 59^. After digestion with Type II collagenase (2mg/mL in PBS/2% BSA, Life Technologies Inc., Carlsbad, CA, USA) at 37°C for 60 min, samples were strained (100uM filters; Corning Inc./Falcon), centrifuged at 250rcf for 10 min, resuspended in RBC, centrifuged and resuspended in preadipocyte medium [Dulbecco’s Modified Eagle Medium/Nutrient Mixture F-12 (DMEM/F12 50:50; Thermo F, Cat# 11330032), 15% fetal bovine serum (FBS; ThermoFisher Scientific Inc./Gibco, Cat# 26140) and 1% Antibiotic-Antimycotic 1% antimycotic-antibiotic solution (ThermoFisher Scientific Inc./Gibco, Cat# 15240062)]. FACS-isolated ASC and bulk ASC were plated at 3x 10 ^4^ cells/cm^2^ density in preadipocyte medium and adherent cells were passaged three times to enrich for preadipocytes.

Adipogenic differentiation of ASC was induced during 14 days with human adipocyte differentiation medium [DMEM/F12 50:50 and the following reagents from Sigma-Aldrich Inc.: 10 mg/l transferrin (Cat# 10652202001), 33 µM biotin (Cat# B0301), 0.5µM human insulin (Cat# 91077 C), 17 µM pantothenate (Cat# C8731), 100nM dexamethasone (Cat# D9184), 2nM 3,3ʹ,5-Triiodo-L-thyronine sodium salt (Cat# T6397), 1µM ciglitazone (Cat# 230950), 540 µM Isobutyl-1-methylxanthine (Cat# I7018), as previously described. ^60^ Minimal differentiation medium was identical to human adipocyte differentiation medium except that it contained reduced insulin (10nM) and no ciglitazone.

### Adipogenesis assays

We evaluated adipogenic capacity of ASC subpopulations *in vitro* by quantifying adipocyte lipid accumulation using AdipoRed Assay, ORO, and LipidTOX staining.^61^ AdipoRed Adipogenesis Assay (Lonza Group, Basel, Switzerland, Cat# PT-7009) was performed by washing medium off with PBS, staining cells with lipophilic AdipoRed^TM^ Assay Reagent, which specifically partitions into the lipid droplets, and measuring fluorescence at 572 nm. For ORO staining, adipocytes were fixed with 4% formaldehyde, stained with 0.2% ORO solution (Sigma Aldrich Inc., Cat# O0625) in 40% 2-propanol for30min, and absorption of eluate measured photometrically at 510nm. ^61^ Lipid measurements were standardized by total protein as measured by Pierce BCA assay (ThermoFisher Scientific Inc., Cat# 23225). HCS LipidTOX™ staining (Thermo Fisher Scientific Inc./Invitrogen; #H344-75/76) was performed in 4% formaldehyde-fixed adipocytes (20 min, RT) using a 1:200 stain:1X PBS dilution (30 min, RT). Cells’ nuclei were then DAPI stained (1 ug/mL; 5 min, RT) and adipocytes imaged with a 20X objective using a Nikon Eclipse Ti inverted point-scanning confocal microscope (Nikon, Tokyo, Japan) equipped with two standard PMT.

### Lipolysis assay

ASC were seeded at 2 x 10^4^ cells/well in 96-well plates and differentiated for 14 days. Before lipolysis induction, adipocytes were washed twice and incubated for 4h at 37°C with warm 0.1% BSA phenol red free DMEM:F12. Lipolysis was induced by incubating adipocytes with 100 uM forskolin (Cayman Chemical Co., Ann Arbor, MI, USA Cat# 11018) for 1h at 37°C in 3% BSA KRBH (135 mM NaCl, 5mM KCl, 1 mM MgSO4, 0.4 mM K2HPO4, 1 mM CaCl2, 5.5 mM Glucose, 20 mM HEPES; pH 7.4). Supernatants were collected for glycerol concentrations analysis (Glycerol Assay Kit, Sigma-Aldrich inc., Cat# MAK117), which were calibrated by cell DNA content (CyQUANT, ThermoFisher Scientific Inc., Cat# C7026). Results are shown in glycerol release fold change of forskolin-stimulated cells over basal conditions (non-stimulated cells).

### Quantitative real time PCR (qRTPCR)

ASC collected immediately after FACS sorting were pelleted, then resuspended in RNA lysis buffer (RLT buffer, Qiagen, Venlo, Netherlands). In vitro differentiated adipocytes were washed twice with cold 1X PBS and collected in RLT buffer. Samples were snap frozen in liquid nitrogen. RNA fwas isolated with RNeasy Mini Kit (Qiagen, Cat# 74104). Purity, concentration, and integrity of RNA were evaluated using a NanoDrop 1000 spectrophotometer (ThermoFisher Scientific Inc.). Equal amounts of input RNA were reverse-transcribed using Applied Biosystems High Capacity cDNA Archive Kit (Applied Biosystems Inc., Foster City, CA, USA). qRTPCR was conducted with TaqMan primers and reagents (Life Technologies Inc., Carlsbad, CA, USA). Data are presented as fold changes calculated from least squares mean differences according to the 2^−ΔΔCt^ method ^62^ and normalized to B2M, or the average expression of B2M and ACTB housekeeping gene controls. Samples used as calibrator are described in each figure.

### Immunofluorescence microscopy

Whole AT samples and cells were fixed, blocked, and permeabilized as described by our group^17^ . Briefly, samples were fixed for 20min in 4% PFA (RT, dark room) and blocked in 1X PBS with 5% BSA + 0.1% saponin (500uL each well) 60 min RT. After 3 consecutive washes with 1X PBS (10 min each), antibody incubations were performed. Lumican primary antibody (ThermoFisher Scientific Inc., # PA5- 14571) was used at 1:100 dilution for 1h, RT in blocking buffer. TM4SF1 primary antibody (Abcam Inc, Cat# ab113504) was used at 1:50 overnight at 4C. Either Alexa 568 (ThermoFisher Scientific Inc.) or Alexa 488 (ThermoFisher Scientific Inc.) at 1:250 dilution for whole tissue and 1:1000 dilution for cells were used for 1h at RT. followed by incubation with Alexa 568 (ThermoFisher Scientific Inc., 1:250) for 1h, RT. Samples were then stained with HCS LipidTOX™ Green or Red Neutral Lipid Stain (1:200; ThermoFisher Scientific Inc., Cat# H34475) for 30 min at RT followed by DAPI staining (1 ug/mL, 5 min, RT). A Nikon Eclipse Ti inverted point-scanning confocal microscope (Nikon, Tokyo, Japan) equipped with two standard PMT detectors was used for obtaining images.

## ACKNOWLEDGMENTS

The University of Michigan Advanced Genomics Core Facility performed RNA sequencing. This work supported by NIH grants R01DK115190 (RWO, CNL), R01DK090262 (CNL), Veterans Affairs Grant I01CX001811 (RWO).

## AUTHOR CONTRIBUTIONS

Conceptualization: RWO, CNL, CSB; Data curation: CSB, JW, RWO, CNL; Investigation: CSB, LG, JD, CGF, TE, OA, AK, APE; Project administration: RWO, CNL; Resources: RWO, CNL; Methodology: CSB, LG, JD, CGF, TE, OA, AK, APE; Funding acquisition: RWO, CNL; Writing, original draft: CSB, RWO, CNL; Writing, review & editing: all authors.

## COMPETING INTERESTS

The authors declare no competing interests.

## Notes

### Competing Interest Statement

The authors have declared no competing interest.

